# The role of genetic constraints and social environment in explaining female extra-pair mating

**DOI:** 10.1101/588475

**Authors:** Daiping Wang, Wolfgang Forstmeier, Katrin Martin, Alastair Wilson, Bart Kempenaers

**Author notes:** Address for correspondence: Wolfgang Forstmeier, Department of Behavioural Ecology and Evolutionary Genetics, Max Planck Institute for Ornithology, Eberhard-Gwinner-Str. 7, 82319 Seewiesen, Germany.

## Abstract

Why females of socially monogamous species copulate with males other than their partner has been a long-standing, unresolved puzzle. We previously reported that female promiscuity appears to be a genetic corollary of male promiscuity (intersexual pleiotropy hypothesis). Here we put this earlier finding to a critical test using the same population of zebra finches *Taeniopygia guttata*. After three generations of artificial selection on male courtship rate, a correlate of extra-pair mating, we assess whether female promiscuity changed by indirect selection and we re-examine the crucial genetic correlations. Our new analyses with substantially increased statistical power clearly reject the hypothesis that male and female promiscuity are genetically homologous traits. Our study highlights that individual females show low repeatability in extra-pair mating behavior across different social environments. This emphasizes the potential importance of pair bond strength and the availability of favored extra-pair males as factors explaining variation in patterns of female promiscuity.

## Introduction

Why females in socially monogamous species actively engage in matings outside the pair bond is a long-standing, intriguing question [1–5]. Mating outside the pair bond is obviously adaptive for males (i.e. benefits from this behavior will typically outweigh costs), because it leads to additional offspring that are raised by another pair, and hence directly increases male fitness [6, 7]. However, why females engage in extra-pair copulations is more puzzling: promiscuous behavior does not increase the number of offspring females can produce and is associated with costs such as increased predation risk, increased risk of contracting sexually transmitted diseases, reduced paternal care and punishment by the social mate [1, 4]. In birds, more than 90% of species breed in socially monogamous pairs, but extra-pair paternity is common [2, 8]. Birds have served as paragons for studying the evolution of female promiscuity, because males typically cannot force copulations and females often actively seek extra-pair copulations [9–11]. The majority of studies tried to explain the occurrence of female extra-pair mating behavior by highlighting the potential benefits [1, 2, 12]. These included indirect genetic [13–15] as well as direct ecological benefits [8, 16, 17]. Yet, despite much empirical work, the general support for these adaptive scenarios remains limited [4, 18–21]. Therefore, alternative, non-adaptive explanations deserve attention [12].

Several hypotheses of ‘genetic constraint’ have been proposed to solve the evolutionary puzzle of apparent non-adaptive female extra-pair behavior [22, 23]. These hypotheses assume that promiscuous behavior is heritable and state that the alleles underlying female promiscuity are maintained in the population, because they have additional pleiotropic effects that are beneficial to at least one sex. Depending on whether the pleiotropic effect is expressed in males or females, two types of hypotheses can be distinguished.

(1) The hypothesis of ‘intersexual pleiotropy’ proposes that female and male promiscuity are homologous traits that are affected by the same sets of genes [22]. Alleles that increase promiscuity will be maintained in the population due to positive selection in males. When inherited to a daughter, these alleles will cause female promiscuity even if this behavior is not adaptive for females. This hypothesis requires a positive genetic correlation between measures of female and male promiscuity (i.e. positive cross-sex genetic covariance).

(2) The hypothesis of ‘intrasexual pleiotropy’ posits that female promiscuity is maintained because its causal alleles have pleiotropic effects on other female traits that are under positive selection [11, 23]. For example, female responsiveness to male courtship might be genetically linked to female fecundity, because courtship may proximately stimulate egg production [24]. Alternatively, genetic variants underlying increased female sexual responsiveness towards her social mate may be favored by selection because low responsiveness can lead to infertility and hence reduced fitness [23]. Positive selection on alleles for increased responsiveness towards the social mate could then lead to increased female responsiveness towards extra-pair males as well. This hypothesis requires that female promiscuity is positively genetically correlated to either female fecundity or to female responsiveness towards her social mate (i.e. within-sex genetic covariance).

Empirical testing of these hypotheses using field data on extra-pair paternity is difficult, because heritability of male and female promiscuity is low [25–28]. The main problem is that the realized patterns of paternity also depend on factors other than the intrinsic inclination of an individual to seek extra-pair copulations, such as sperm competition and mate guarding.

In an earlier study on captive zebra finches [29], we combined data on realized levels of extra-pair paternity with detailed observations on behaviors that reflect an individual’s propensity to engage in extra-pair mating. We found strong, positive genetic correlations between male and female measures of extra-pair mating behavior, supporting the ‘intersexual pleiotropy’ hypothesis. We rejected the ‘intrasexual pleiotropy’ hypothesis, because the genetic correlation between responsiveness to the partner and responsiveness to extra-pair males did not differ from zero. Our study thus suggested that female promiscuity can be changed indirectly by artificially selecting males for increased or reduced courtship rate, a genetic correlate of male extra-pair siring success and of female promiscuity.

The present study reports on the results of such an artificial selection experiment. Using the birds from the initial study, we set up two replicate lines for high male courtship rate, two replicate lines for low courtship rate and two unselected control lines. Increasing the genetic variance in male courtship rate allowed us to test with increased statistical power whether female extra-pair mating behavior is indeed genetically linked to male courtship rate. Based on our previous results, we predicted that the level of female promiscuity would change indirectly by selection imposed on male behavior only.

This study also amends a weakness of the initial study: previously, we measured the behavior of a female only once, in the context of being paired to the partner she had chosen in an experiment. The observed behavior was then assumed to be representative for that female. However, extra-pair behavior might also have been a property of the female’s social environment (e.g. strength of the social pair bond, characteristics of the available extra-pair males). To resolve this, we here measured extra-pair behavior of each female with two successive partners. This allows to assess the amount of variation that is due to the female and to the environment. Thus, we examine the repeatability of female promiscuity and quantify its heritability and genetic covariance with other traits.

To examine the ‘intersexual pleiotropy’ hypothesis, we quantified the sign and strength of the genetic correlations between measures of female promiscuity and two measures of male sexual behavior, namely (1) male courtship rate (under artificial selection), and (2) male success in siring extra-pair eggs. To test the ‘intrasexual pleiotropy’ hypothesis, we quantified the correlations between female promiscuity and (3) female responsiveness towards her social mate, and (4) measures of total female fecundity.

## Results

### Selection lines for male courtship rate

We established six selection lines and bred them over three consecutive generations: two lines selected for high male courtship rate, two for low courtship rate, and two unselected control lines. Figure 1 shows, for each generation, the actual phenotypes (courtship rate) of all male offspring that were bred as a function of the mean breeding value of their parents (i.e. as a function of the predicted offspring phenotypes based on a genetic model that includes the observed phenotypes of parents and their relatives). The slope of the regression lines is close to unity, indicating that the offspring generations behaved as predicted by the genetic model. With each generation, we chose parents with even more extreme breeding values, as reflected by the outward movement of the high and low lines along the x-axis over progressive generations (Figure 1A-C). In consequence, the offspring phenotypes became progressively differentiated along the y-axis between the selection lines. After three generations of selection, the average difference between the high and the low lines reached 2.4 phenotypic standard deviations (Cohen’s d; [30]). The two replicates of each type of line behaved almost identically (Figure 1, Table S3).

**Figure 1.**
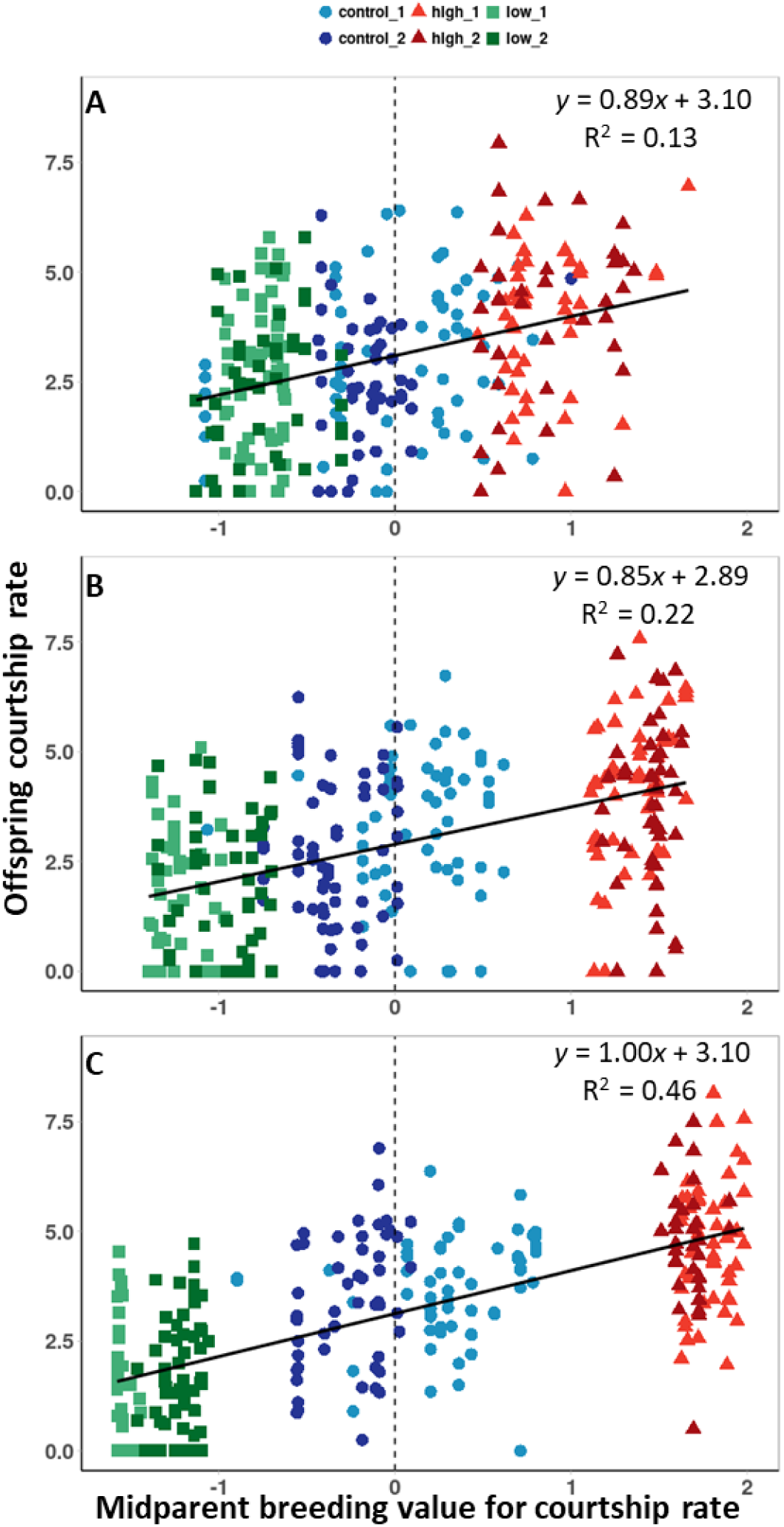
Male observed and predicted courtship rate in individuals from six selection lines over three successive generations (S1-S3, A to C). The y-axis shows the measured courtship rate of male offspring (seconds in a 5-min trial, averaged across 4 trials per male, square-root transformed). The x-axis shows the predicted courtship rate, that is, the parents’ breeding value for male courtship rate. These values were estimated prior to breeding (generations S0-S2, without information on offspring phenotypes) from a single-trait permanent-environment animal model in VCE. Symbol color and shape indicate the three types of selection lines (high, control, and low). Within each type, light and dark colors indicate the two replicate lines. Ordinary least-square regression lines and their equations are shown.

### Indirect response to selection

We assessed whether the successful selection on male courtship rate resulted in correlated changes in levels of extra-pair paternity in both sexes. To this end, we put equal numbers of males and females from the three types of selection lines in communal aviaries, noted pair formation and let the birds breed for two breeding rounds, each lasting seven weeks during which females laid up to three clutches. We then quantified for each individual the level of extra-pair paternity.

During the time they were monogamously paired, 190 females produced 2,951 fertile eggs, 726 of which (24.6%) were sired by extra-pair males. Levels of extra-pair paternity (% of extra-pair young in all broods) ranged from on average 37.4% in line 1 ‘high’ to 15.8% in line 2 ‘low’, with the other four lines showing intermediate levels (Figure 2). An analysis of individual levels of extra-pair paternity with selection regime (coded as a continuous variable: 1df; low = −1, control = 0, high = 1) as the predictor of interest, showed a significant effect (β = 0.698, z = 3.1, p = 0.002, n = 190, Table S4).

**Figure 2.**
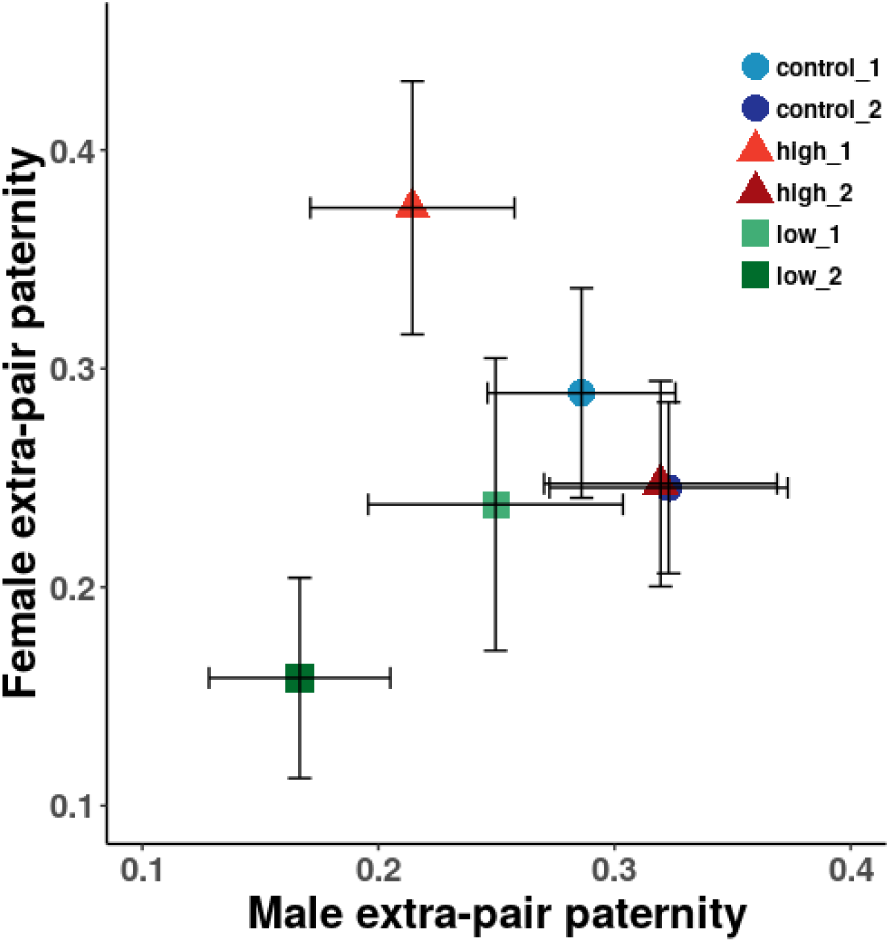
Weighted averages (± SE) of levels of male and female extra-pair paternity for each of the six selection lines in aviary breeding experiments (data on ca. 3,000 eggs from the ‘S3’ generation, see Results). Male extra-pair paternity (x-axis) refers to the total proportion of eggs sired by socially paired males outside their pair bond. Female extra-pair paternity (y-axis) refers to the proportion of eggs laid by socially paired females that are sired by males other than the social partner.

From the male perspective, 188 individuals sired 3,067 eggs during the time they were socially paired, 851 of which (27.7%) with females other than their social mate. The corresponding average levels of extra-pair paternity (% of all young sired with extra-pair females) ranged from 32.2% in line 2 ‘control’ to 16.7% in line 2 ‘low’ (Figure 2). Here, selection regime showed a non-significant trend in the expected direction (β = 0.278, z = 1.7, p = 0.09, n = 188, Table S5).

### Repeatability of female promiscuity

After one round of breeding and a break of two weeks (housing in unisex groups), we rearranged all individuals for a second breeding round. Each individual was then allowed to breed again for seven weeks with a different social mate and a different set of potential extra-pair mates. Estimates of repeatability for the female’s responsiveness to courtship by extra-pair males (‘female extra-pair response’) and for the level of extra-pair paternity are relatively low (Figure 3A, B). This is confirmed by animal models showing that the random effect of social pair (‘Pair ID’) explained considerably more variance in measures of female promiscuity than the random effects that represent female identity (Figure 3C; animal models of ‘Genetic’ + ‘Permanent environment’: Tables S6, S7, S10 to S15). In other words, a female’s level of promiscuity is more consistent within a given context (social pair bond, set of extra-pair males) than between contexts (Figure 3C).

**Figure 3.**
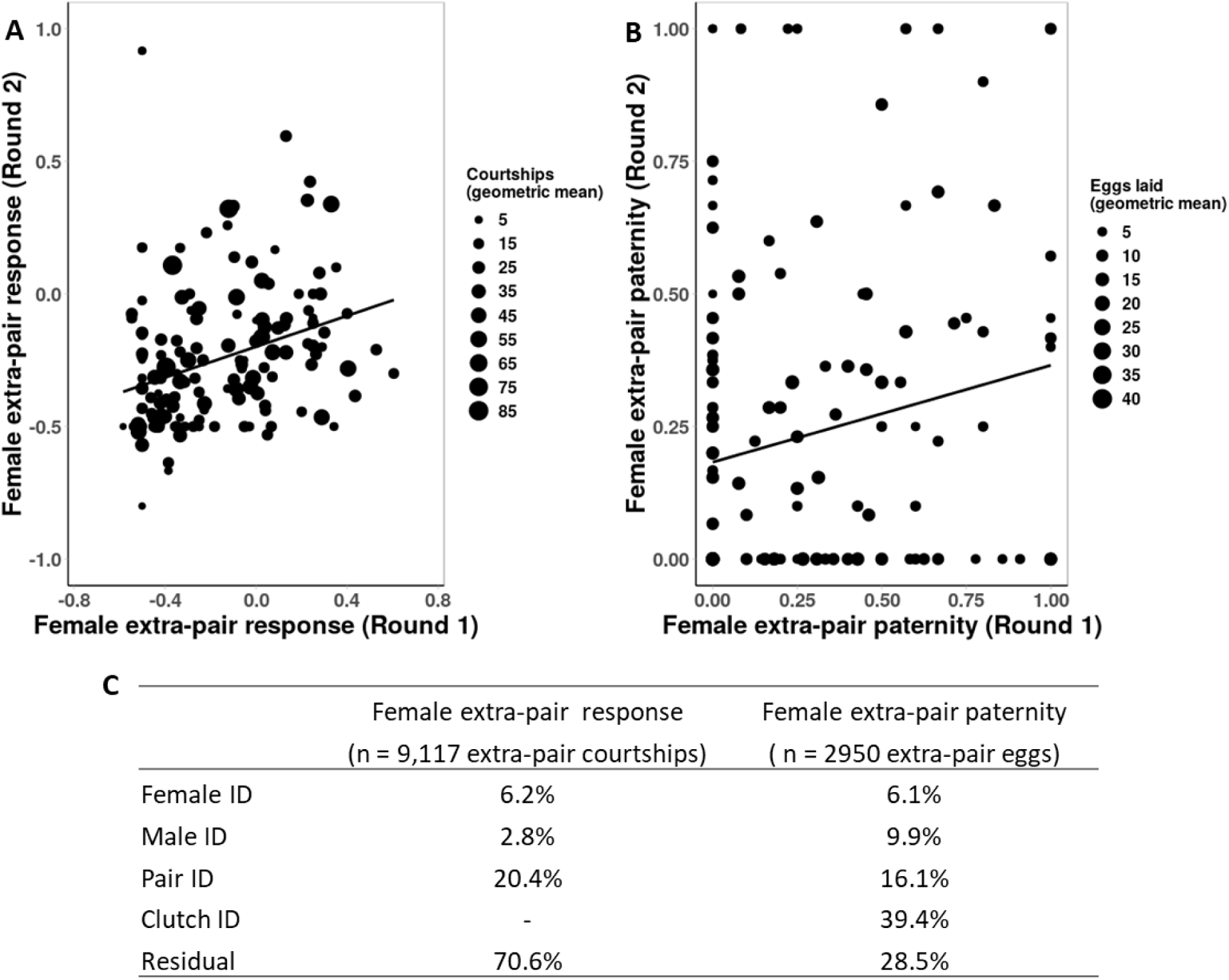
Estimates of repeatability of ‘female extra-pair response’ (A, the responsiveness of females to courtship by extra-pair males) and levels of extra-pair paternity (B, proportion of eggs sired by extra-pair males) across two breeding rounds differing in social pair bonds and in the identity of potential extra-pair males. Shown are ordinary least square regression lines weighted by the geometric mean of the two rounds ((A) slope β = 0.37 ± 0.09, N = 151 females; (B) β = 0.24 ± 0.09, n = 135 females). Dot size refers to the geometric mean of the relevant sample sizes in the two breeding rounds (number of extra-pair courtships and number of eggs laid, respectively). (C) Variance components estimation of the random effects based on mixed-effect models with ‘female extra-pair response’ and ‘female EPP’ (each egg modeled as 0 = within-pair and 1 = extra-pair) as the dependent variable, respectively.

### Testing the ‘intersexual pleiotropy’ hypothesis

Figure 4 shows the relationship between measures of female extra-pair behavior and the female breeding values for male courtship rate. All slopes are positive, but slopes based on data from the initial study (Figure 4A, B) are steeper than those based on data from the selection lines (Figure 4C, D), whereby the latter are the more powerful tests. The genetic correlations between measures of male and female promiscuity are presented in Figure 5; estimates of between-sex genetic correlations from 5-trait animal models based on the initial data (Figure 5A Table S10, S11) are contrasted with estimates from models based on data from the lines artificially selected for high and low male courtship rate (Figure 5B; Tables S12, S13). The latter show between-sex genetic correlations close to zero for male courtship rate (median of four estimates: r_A_= 0.04, Figure 5B; Table S18), and negative values (i.e. opposite to expectations) for male extra-pair siring success (median r_A_ = −0.34, Figure 5B; Table S18). These estimates stand in strong contrast to the positive estimates derived from the initial data (Figure 5A). An updated matrix of genetic correlations estimated from the joint data (initial plus selection lines) shows weekly positive genetic correlations that are not significantly different from zero (Figure 5C; summary of Tables S6 to S9 showing medians of estimates from four types of models).

**Figure 4.**
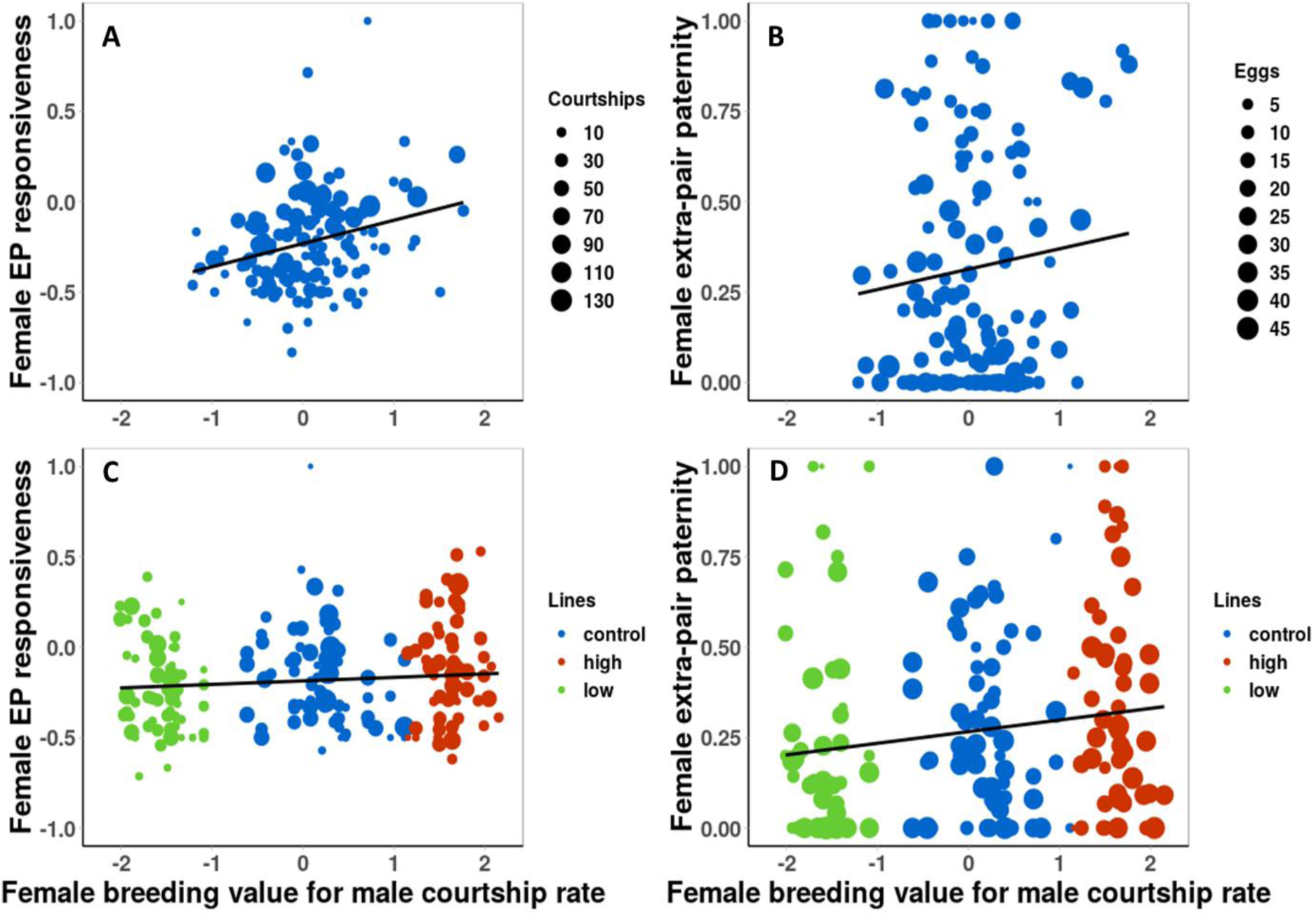
Relationships between measures of female extra-pair behavior and their estimated breeding value for male courtship rate. (A) and (B) Data from Forstmeier et al. (2011) (generations F1-F3)[29]; (C) and (D) data from the selection lines (generation S3). (A) Average female responsiveness to courtship by extra-pair males (‘Female EP responsiveness’, n = 141 females, 3,958 courtships) in relation to their estimated breeding value for male courtship rate. Dot size refers to the number of extra-pair courtships observed for each female (range: 1–138, median: 19). Shown is the regression line weighted by the number of courtships (slope β = 0.14 ± 0.03). (B) Average level of female extra-pair paternity (the proportion of eggs sired by extra-pair males; n = 149 females, 2,253 eggs) in relation to their estimated breeding value for male courtship rate. Dot size refers to the number of eggs laid by each female (range: 1–45, median: 14). Shown is regression line weighted by the number of eggs (β = 0.10 ± 0.04). (C) Average female responsiveness to extra-pair male courtship (n = 205 females, 9,117 courtships) in relation to their estimated breeding value for male courtship rate. Colors refer to the type of selection line (control, high, low). Dot size refers to the number of extra-pair courtships observed for each female (range: 1–219, median: 33). Shown is the weighted regression line (β = 0.02 ± 0.01). (D) Average level of female extra-pair paternity (n = 190 females, 2,951 eggs) in relation to their estimated breeding value for male courtship rate. Dot size refers to the number of eggs laid by each female (range: 1–32, median: 15). Shown is the weighted regression line (β = 0.03 ± 0.01). Female breeding values for male courtship rate come from a single-trait permanent environment model conducted in VCE based on courtship rates from 800 (A,B) and 1,651 (C,D) male relatives. Note that the regression lines are for illustration only, because other influential fixed effects are not taken into account.

**Figure 5.**
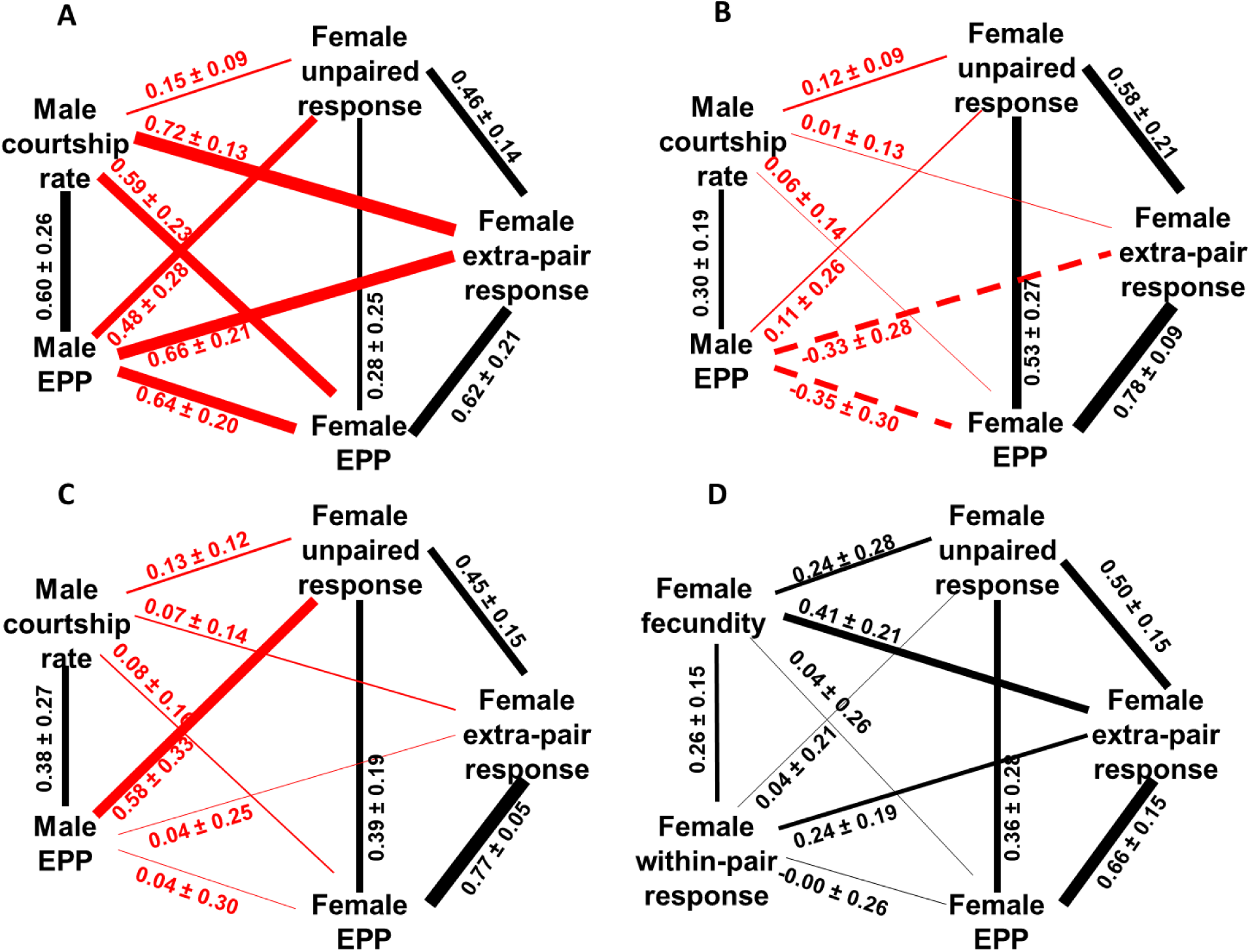
Estimates of genetic correlations between measures of male and female extra-pair mating behavior. A, initial data from the previous study (29). Shown are median estimates (± median SE) from two versions of animal models (Table S10 and S11; see description of models 5 and 6 in Methods). B, selection lines. Shown are median estimates (± median SE) from two versions of animal models in this study (Table S12 and S13; see description of models 7 and 8 in Methods). C, joint data from the initial study and the selection lines. Shown are median estimates (± median SE) from four versions of animal models (Table S6 to S9; models 1 to 4 in Methods). D, genetic correlations among female traits. Shown are median estimates (± median SE) from four versions of animal models (Table S14 to S17; models 9 to 12 in Methods). Between-sex genetic correlations are shown in red, within-sex genetic correlations in black. Line thickness reflects the strength of the correlation. ‘Female EPP’: paternity of each egg laid by a paired female, scored as 0 = within-pair and 1 = extra-pair; ‘Male EPP’: the number of extra-pair eggs males sired within each breeding round. Note that all traits are measured during breeding in communal aviaries except for “male courtship rate” and “female unpaired response” which reflect the behavior of unpaired birds in standardized cage trials (see Methods).

### Testing the ‘intrasexual pleiotropy’ hypothesis

We found a moderately strong positive genetic correlation between female responsiveness to extra-pair male courtship (‘female extra-pair response’) and female fecundity (Figure 5D), yet its estimated strength varied considerably across different models (between 0.05 and 0.59, Table S14 to S17). Estimated genetic correlations between female extra-pair and within-pair response were weakly positive (Figure 5D), but also not robust (see Tables S14 to S17). Note that genetic correlations involving ‘female within-pair response’ are particularly difficult to estimate because the trait shows low repeatability across pair bonds (see Methods: Data Analysis: Female extra-pair and within-pair response).

## Discussion

Overall, our data show high context-dependence of female promiscuity and more support for the ‘intrasexual pleiotropy’ hypothesis than for the hypothesis of ‘intersexual pleiotropy’. This study thus suggests that female promiscuity is an ‘independent trait’ of females rather than a ‘corollary’ of male promiscuity [31, 32].

The breeding of selection lines for male courtship rate was effective in maximizing the statistical power for testing whether measures of female promiscuity are genetically correlated with male courtship rate as a proxy of male promiscuity (see the increased data range in Figure 1 and Figure 4). Based on the most decisive test for such a genetic correlation (see ‘new data’ in Figure 5B; Table S18), we reject the ‘intersexual pleiotropy’ hypothesis, despite weak supportive trends in the phenotypic data (Figure 2 and Figure 4D) and weak, positive correlations in the analysis of the joint data (Figure 5C). Statistical testing suggested a significant effect of the selection regime on female levels of extra-pair paternity, mostly stemming from reduced levels of extra-pair paternity in females from the two lines for low male courtship rate (Figure 2, Table S4). However, we base our conclusions on animal models that control for non-independence of individuals in the different selection lines via genetic relatedness (Figure 5).

We found a significant, positive genetic covariance between female responsiveness to extra-pair males (‘female extra-pair response’) and female fecundity, and a somewhat lower positive genetic covariance between female extra-pair responsiveness and her responsiveness to her social mate (‘female within-pair response’) (Figure 5D). This finding should be interpreted cautiously (given that the estimates did not seem robust, see also below) and deserves more study, in particular from populations of different species breeding in the wild.

Our study reveals strong context dependence of female extra-pair mating behavior: ‘Pair ID’ explained more variation than female identity (Figure 3, Tables S6, S7). This could reflect variation in the quality of the social pair bond, or in the set of available extra-pair males, which can be studied further under a social network framework [5].

### Comparison of the initial study with this study

The conclusions from this study and from our earlier work [29] differ substantially. We discuss several potential explanations for this difference. First, the initial study was based on a smaller sample of 150 females. Hence, founder effects [33] may have resulted in some linkage disequilibrium between alleles for male and female promiscuity by chance alone. Such non-physical linkage may then have broken up during the breeding of selection lines. Second, the measures of female extra-pair behavior (mean phenotypes; y-axes in Figure 4) are noisier in the initial study than in this study, because in the latter they are based on two rounds of breeding with different social mates and a different set of potential extra-pair partners. Third, estimates of female breeding values for male courtship rate (x-axes in Figure 4) were based on half the number of male relatives in the initial study compared to this study, leading to higher error along the x-axis in the initial study. Updating the breeding value estimates of the females involved in the initial study with the new information on courtship rate of their sons, grandsons and great-grandsons, already leads to considerably shallower regression slopes in Figure 4A (β = 0.14) and 4B (β = 0.10).

In conclusion, we suggest that the significant finding in our initial study [29] is a type I error resulting from relatively noisy data. There is no evidence that inadequate modelling caused the difference, because updating the earlier models by including clutch and pair identity (‘Clutch ID’, ‘Pair ID’) as additional random effects, did not alter the conclusions (see Figure 5A and Tables S10 and S11). Note that estimates from Bayesian models in MCMCglmm were smaller, had larger standard errors and were closer to estimates from the follow-up study than those from REML models in VCE (Table S18). This confirms the notion that the estimation of genetic correlations can be problematic when heritabilities are relatively low [34, 35] and sample sizes are limited.

### Future directions and conclusions

Female fecundity was positively genetically correlated with measures of female promiscuity (Figure 5D), but Bayesian models in MCMCglmm again yielded more conservative estimates (median r_A_ = 0.14 ± 0.21) than REML models in VCE (median r_A_ = 0.59 ± 0.20). To assess whether genetic covariance with fecundity is a more general explanation for the persistence of female extra-pair mating, follow-up studies in the wild will be needed. Reid et al. [27] reported positive genetic covariance between female levels of extra-pair paternity and female annual reproductive success, but it is unclear whether this was due to variation in fecundity or variation in rearing success. If quantitative genetic analyses are not feasible because detailed pedigree information is not available, one could still examine whether there is a positive phenotypic correlation between clutch size and levels of extra-pair paternity. Such analyses, however, would need to take into account the mechanisms behind extra-pair paternity. For example, the probability of detecting extra-pair paternity (i.e. that an extra-pair copulation leads to a fertilization) might increase with clutch size. In field studies, it may also be important to control for breeding density, because the latter may influence both the availability of extra-pair males and clutch size.

Our analyses of female extra-pair behavior across two social environments (Figure 3) revealed a substantial amount of context-dependence of this behavior. When considering levels of extra-pair paternity, the most influential factor was the identity of the social pair (‘Pair ID’, Figure 3C), indicating consistency across multiple clutches with the same partner and flexibility in behavior when breeding with different partners (Figure 3B). Such consistency at the level of the social pair rather than at the level of the female (‘Female ID’, Figure 3C) is consistent with findings in coal tits [36] and suggests that extra-pair paternity levels may vary with the strength of the social pair bond (e.g. behavioral compatibility of mates, as suggested by [37]). Similarly, a female’s responsiveness to courtship of extra-pair males strongly depended on the combination of male and female identities, i.e. on who courted whom (coded as ‘Pair ID’ in Figure 3C; Tables S6, S7). Hence, the occurrence of promiscuous behavior may depend more strongly on aspects of compatibility between individuals. The dependence on the social context might reflect the quality of the social pair bond or the availability of specific extra-pair males, or both. The relative importance of these factors could be addressed by targeted experiments or by social network analyses [5].

Contrary to our previous claim, the artificial selection experiment showed that levels of female promiscuity cannot be altered by artificially selecting on the courtship rate of unpaired males (a correlate of extra-pair siring success that can be measured prior to pairing), at least not within the range covered by our selection lines. Nevertheless, this study suggests that models of genetic constraint remain in general a viable explanation for the persistence of female extra-pair mating. All examined genetic correlations in Figure 5 (A and B) were positive (instead of 50% as expected from randomness). Examining these constraints in other study systems appears both promising and feasible.

## Methods

### Subjects

All study subjects come from a population of zebra finches that has been maintained at the Max Planck Institute for Ornithology in Seewiesen, Germany since 2004 (population # 18 in [38]). Housing conditions, diet and aviary specifications for breeding have been described in detail in the supplementary file to [39]. For this study, the pedigree of this population comprises eight generations: Parental, F1 to F4, and four generations of selection lines (S1 to S3, see below).

### Behavioral Observations

We measured behavioral traits related to extra-pair mating under two experimental set-ups: (1) in cages, where behavior could be measured under standardized conditions, leading to high individual repeatability; (2) in aviaries, where individuals bred repeatedly and were exposed to different sets of potential extra-pair partners.

#### a) Cage Experiments on Unpaired Birds

Before the formation of social pair bonds, we measured for each male in the population ‘male courtship rate’ (the trait subjected to artificial selection) towards an unpaired female introduced into his cage. We set up encounters between an unpaired male and an unpaired female that were unfamiliar to each other. Each encounter (‘trial’) lasted five minutes during which we recorded the total duration (in seconds) of male courtship, that is, song directed towards the female. For each female, we scored her responsiveness to the male (‘female unpaired response’) during each encounter on a five-point scale following [11], where −1 represents a clear rejection (involving aggression, threat, or fleeing) and +1 a clear acceptance (involving copulation solicitation, beak wiping, and ritualized hopping) with intermediate scores (−0.5, 0, +0.5) given for weaker or mixed responses [11, 29]. For this study, we combined 3,776 trials from the initial study [29] and 3,014 trials on individuals from the selection lines (see below). In total, we obtained 6,786 measures of ‘male courtship rate’ (four encounters with missing data were excluded) and 5,039 measures of ‘female unpaired response’ (74% of all trails; responsiveness could not be scored in 1,751 trials, typically when there was no male display). The trials involved 1,556 males and 1,441 females and were carried out between July 2002 and December 2013. Males encountered on average 4.4 ± 1.3 SD (range 2-8) different females, and females encountered on average 4.5 ± 2.2 SD (range 1-14) different males (Table S1).

##### Selection on Male Courtship Rate

We established lines selected for divergent breeding values for male courtship rate, starting in 2009 [some details see 40, 41].

##### Founder generation ‘S0’

Before initiating the breeding of selection lines, we measured the courtship rate of 585 males from four consecutive generations (P to F3, not including F4 birds) [29] in 2,922 trials. Using these measurements, we estimated breeding values for male courtship rate with a pedigree-based animal model. Breeding values of all individuals in the pedigree (n = 1219 from P to F3, including females) were calculated using VCE 6.0.2 [42]. The single-trait permanent-environment animal-model was set up as follows. (1) ‘Male courtship rate’ was squared-root transformed to approach normality and used as the response variable (Table S1). (2) Fixed effects were male test day (four levels, from day one to day four), time of day of the trial start (continuous, range: 8:51-18:19), male inbreeding coefficient F (continuous, range: 0-0.25) and rearing environment of the male (two levels, mixed-sex or unisex). (3) As random effects we included ‘Animal’ (additive genetic effect), ‘Male ID’ (permanent environment effect, 585 levels), ‘Female ID’ (maternal effect, 203 levels), ‘Test batch ID’ (period of testing, 8 levels), and ‘Cohort ID’ (periods of breeding, 6 levels).

We started six breeding lines (two control, two high and two low lines) by choosing founder individuals with the estimated breeding values for courtship rate (see above) from the pool that were still alive in May 2009 (n = 773; see Table S19). For each line, we let 15 pairs breed in one of 90 randomly assigned cages (60×40×45cm) distributed over two breeding rooms (45 cages each). First, we randomly selected birds from the entire pool for the two control lines. Then, we selected 30 birds of each sex with the highest breeding values for courtship rate for the two ‘high’ lines, and randomly allocated half of them to each replicate line. Thereafter, we also selected six ‘replacement’ individuals of each sex (in case a high line bird would die during breeding) with the next highest breeding values and distributed them randomly among the two lines. The two low lines were selected in the same manner, but using the birds with the lowest breeding values.

Within each line, the 15 breeding pairs were chosen in such a way as to minimize the level of inbreeding (see Table S19). Each pair was allowed to breed in two ‘rounds’ over a total period of about 14 months (from pair formation to independence of the last offspring). In each round, we allowed pairs to breed until we obtained about 50 juveniles from each line. After round one, we redistributed the birds within each line such that they obtained a new partner (breeding cages again randomly assigned). In this way, we created maternal and paternal half sibs, which facilitated the separation of maternal effects from additive genetic effects. We placed juveniles (age: 35 to about 120 days) of each breeding round in one of two large, mixed-sex groups. Thus, across both rounds of breeding of each generation (S0, S1 and S2, details see below), roughly 600 offspring were raised in four mixed-sex groups comprising roughly 75 males and 75 females from all lines.

##### Breeding generations ‘S1’ to ‘S3’

Birds of the S0 generation produced 568 offspring of which 546 survived until we started breeding the next generation (see Table S19). ‘Male courtship rate’ and ‘female unpaired response’ of these offspring were measured four times per individual (age of testing is given in Table S19). These new measurements were added to update the animal model (with the same fixed and random effects) for the calculation of predicted breeding values for all individuals (n = 1,929). The new model included 4,362 measurements of courtship rate from 947 males.

We selected the S1 breeders (15 pairs plus five replacement birds of each sex in each line) as described above (random selection for control lines and based on breeding values for high and low lines; Table S19). Again, we assigned breeding pairs in such a way as to minimize and standardize the average inbreeding coefficient. Specifically, in the most inbred line (high 2), we minimized inbreeding, while in the other five lines we chose pairs to match the mean value for this line. The mean inbreeding coefficients of the resulting offspring for each line are given in Table S19. The following generations S2 and S3 were bred following the same principles (see Table S19 for summary statistics).

#### b) Aviary Experiments of ‘S3’ Birds

The S3 generation of the six selection lines consisted of 343 female and 338 male offspring, most of which had been phenotyped for ‘male courtship rate’ and ‘female unpaired response’ in the cage experiments (see Table S19). For a subset of 219 females and 217 males (about equally representing the six lines), we also measured other phenotypes directly linked to extra-pair mating.

Between January 2014 and May 2015, we set up 9 breeding aviaries equipped with cameras as described in [29] and let birds breed, as follows. We created four consecutive testing cohorts, each comprising 54 males and 54 females randomly drawn from the available pool of birds in each line (9 males and 9 females from each line per cohort, 216 of each sex in total, plus a few replacements, see below). Each group was distributed over the nine aviaries such that (1) all birds within an aviary were unfamiliar with each other and (2) each aviary contained one male and one female from each selection line. Due to a shortage of line 1 ‘low’ and later also line 2 ‘high’ birds, we used individuals from line 2 ‘low’ and line 1 ‘high’, respectively, in 11 out of 36 rounds of breeding in aviaries. In all cases, aviaries contained 2 males and 2 females from each line type, but overall the number of tested birds per line and sex varied from 25 to 47 (Table S19).

With this setup, each individual had a choice of 6 potential mates. Social pairing appeared random with regard to line (details not shown). Each set of birds spent seven weeks in the aviary, during which most females laid three clutches; nest boxes were provided from day 1 to day 45. We collected all laid eggs for parentage assignment as soon as we found them and replaced them by plastic eggs. Clutches (of plastic eggs) were removed after 10 days of incubation to encourage the female to lay the next clutch. On day 49, all individuals were separated by sex and placed into different rooms for a two-week period, after which we initiated an identical, second round of breeding with a different set of potential social and extra-pair partners (by swapping the six males of one aviary to the next). This allowed us (a) to quantify the repeatability of the measured traits with different partners, and (b) to disentangle effects of ‘Female ID’ from those of ‘Male ID’ and ‘Pair ID’. In the second round, on average 25% of individuals were familiar to each other due to the joint rearing in one of four large natal groups. Overall, one male and three females died during the first breeding round and they were replaced by an individual from the same line in the second round, leading to a total of 217 males and 219 females participating in the experiments.

We fitted all breeding birds with randomly assigned colored leg bands for individual recognition and observed their behavior. Observations lasted about 30 min (for the 9 aviaries combined) and were carried out about 120 times per breeding round. We recorded all instances of “bonding behavior”: allopreening, sitting in body contact or close to each other, and visiting a nest-box together. The start of a pair bond was defined as the day on which >50% of bonding behaviors were directed to a single male (with a minimum of eight observations on this female-male combination; see [43] for details).

Following the initial study [29], we used video cameras to monitor the birds’ courtship behavior continuously in each aviary. Because courtship was most frequently observed in the early morning, we analyzed the first hour of recording on every day during the breeding period, plus another two randomly selected hours per day. In total, we screened 10,656 hours of video (3h × 49.33 days × 9 aviaries × 2 breeding rounds × 4 testing cohorts) at 8-fold speed (equal numbers of hours randomly allocated to two observers D.W. and K.M.), and detected a total of 33,003 courtships. Of those, we scored ‘female extra-pair response’ based on 9,121 courtships of paired females by potential extra-pair males (involving 206 females) and ‘female within-pair response’ based on 13,268 courtships by the social partner (involving 200 females). For each courtship, a single person (K.M.) scored female responsiveness as in the initial study [29]: threat or aggression toward the male (−1), flying away (−0.5), mixed or ambiguous signs (0), courtship hopping and beak wiping (+0.5), and copulation solicitation (+1).

Data from the initial study consisted of 3,958 scores of ‘female extra-pair response’ (from 141 females) and 4,601 scores of ‘female within-pair response’ (from 143 females; Table S1) [29].

### Paternity Analysis

In total, we collected 4,041 eggs and placed them in an incubator for 4 days to obtain embryonic tissue for parentage analysis. We failed to analyze parentage for 685 eggs (14 eggs without yolk, 24 broken eggs, 632 apparently infertile eggs and 15 lost samples or samples with too low DNA concentration). The remaining 3,356 eggs were unambiguously assigned to parents using 15 microsatellite markers [39], but four eggs were only assigned to their mother (due to parthenogenesis, mosaicism, or siring by sperm from the previous experimental round).

We quantified the proportion of extra-pair young for each female (‘female EPP’) based on a subset of 2,951 eggs laid by paired females (726 eggs were sired by an extra-pair male, 24.6%). Similarly, we quantified male extra-pair siring success (‘male EPP’) as the number of eggs a male sired with a female other than its social mate (3,067 eggs, of which 851 were extra-pair sired, 27.7%; the total number of eggs is higher because it includes paired males siring extra-pair offspring with unpaired females).

Data from the initial study included ‘female EPP’ from 2,253 eggs laid by 149 females and measures of ‘male EPP’ from 152 males (Table S1) [29].

### Female Fecundity

We quantified ‘female fecundity’ as described in [39]. In brief, ‘female fecundity’ is the total number of eggs laid by a female within one breeding round (45 days, see above), determined based on a combination of genetic assignment of maternity (3,356 eggs) and social assignment of eggs that could not be genotyped based on observations of nest attendance (610 eggs). For genotyped eggs, “social assignment” was correct in 93.1% of cases (false assignments resulted from egg dumping or nest take-over; [44]). Thus, assignment errors appear negligible compared to the error when omitting all non-genotyped eggs. In total, we obtained 432 estimates of female fecundity (216 females × 2 breeding rounds, involving 219 individuals) based on 3,966 assigned eggs (mean ± SD = 9.2 ± 5.1, range 0-22). To increase statistical power for quantifying genetic covariance between female fecundity and measures of promiscuity, we included data on female fecundity from seven other aviary breeding experiments with genetic parentage assignment (carried out between 2005 and 2017 and involving 6 generations, the same genetic population as the selection lines). This includes data from the first four breeding experiments used in the initial study [29]. Thus, we used a total of 854 fecundity estimates from 461 individual females based on the assignment of 9,127 eggs (mean ± SD = 10.7 ± 6.8, range 0-38). We statistically accounted for potential differences between the eight breeding experiments (see below).

### Data Analysis

Sample sizes and descriptive statistics of the data used for quantitative genetic analyses are given in Table S1 (including the data from the initial study, [29]). We used similar models as in the initial study, except that we included additional random effects (e.g. ‘Pair ID’ and ‘Clutch ID’) and modelled an effect as random instead of fixed (e.g. ‘Test Batch ID’), where appropriate. To examine whether conclusions of the initial study depended on these decisions about model structure, we repeated the initial analyses with the updated model structure.

#### a) Mixed-effect Models Testing Extra-pair Paternity Levels of the Selection Lines

We tested whether individuals from the high lines had higher levels of extra-pair paternity than those from the low lines after three generations of selection on male courtship rate. We used mixed-effect models in the lme4 package in R 3.4.0 [45, 46] to test for differences in EPP levels across the six selection lines. For each sex, the number of extra-pair eggs of an individual within each round was the dependent variable (binomial model of counts of extra-pair young versus within-pair young using the ‘cbind’ function in R). As the fixed effect of interest, we fitted ‘selection regime’ as a covariate with one degree of freedom (low lines = −1, control lines = 0, and high lines = 1). As random effects, we included either ‘Female ID’ (for female EPP, n = 190) or ‘Male ID’ (for male EPP, n = 188), ‘Selection Line ID’ (six levels), and ‘Individual within breeding round ID’ (each line in the data sheet, n = 325 in females and n = 319 in males as an ‘observation-level random effect’ [47] to control for overdispersion of counts arising from the non-independence of eggs within an individual’s breeding round).

#### b) Statistical Approach for Quantitative Genetic Models

First, we used generalized linear mixed-effect models [45, 46] to investigate how each of the traits measured in this study depended on a range of fixed effects. Details of fixed and random effects given below refer to the joint data set (Table S2: initial study plus data from selection lines).

##### Male courtship rate

‘Male courtship rate’ was square-root transformed to approach normality (Table S1). ‘Male courtship rate’ declined significantly over consecutive test days, declined with time of day, declined with male inbreeding coefficient, and was higher for males from a mixed-sex rearing environment compared with the unisex (Table S2). After accounting for these fixed effects, the random effects ‘Male ID’ and ‘Test Batch ID’ (19 levels) explained 46% and 13% of the variance, respectively.

##### Male EPP

The number of extra-pair eggs males sired within each breeding round (‘Male EPP’) was square-root transformed to approach normality, and was modelled as the dependent variable (Table S1). ‘Male EPP’ increased strongly with the number of days the male was paired. This fixed effect controls for variation in the duration of the breeding period and in the duration of the period a male was unpaired. ‘Male EPP’ also declined with male inbreeding coefficient (Table S2). The random effects ‘Male ID’ and ‘breeding year’ (six levels) explained 21% and 8% of the variance, respectively.

##### Female unpaired response

The responsiveness of unpaired females to male courtship (‘female unpaired response’ in cages) differed significantly among consecutive test days (4 levels) and was higher for females reared in mixed-sex as opposed to unisex groups (Table S2). The random effects ‘Female ID’ and ‘Test Batch ID’ (19 levels) explained 37% and 13% of the variance, respectively.

##### Female extra-pair and within-pair response

Females interacted with an average of 5.5 ± 2.4 different extra-pair males (range 1-12; 97% of 346 females with two or more). ‘Female extra-pair response’ declined strongly with time after sunrise and with the duration of the pair bond (days paired). Based on the initial study [29], we assumed that ‘female extra-pair response’ varied over the fertile cycle with highest responsiveness 3 days before the start of egg laying (day 0) and with a continuous decline over the laying sequence. Hence, the fertile cycle was modeled as the number of days from day −3 (6 levels: from 0 to 5, > 5 also coded as 5). Since 2007, all courtships had been scored by the same observer (K.M.). However, we also used data from two additional observers in 2006, so we included observer ID as a fixed effect. Scores of female extra-pair response varied slightly among the three observers (Table S2). The random effects ‘Female ID’, ‘Pair ID’ (i.e. the combination of identities of the courted female and the courting extra-pair male) and ‘Year’ explained 5%, 23% and 1% of the variance, respectively.

The ‘female within-pair response’ declined strongly with time after sunrise, and increased strongly with the duration of the pair bond (days paired). Within-pair responsiveness varied similarly over the fertile cycle as extra-pair responsiveness (Table S2). The random effects ‘Female ID’, ‘Pair ID’, and ‘Year’ accounted for 1%, 15% and 5% of the variance, respectively.

##### Female EPP

The dependent variable ‘Female EPP’ was modeled for each egg laid by a paired female, as 0 = within-pair and 1 = extra-pair (5,194 eggs in total). This model used a Gaussian error structure, because models with binomial error structure did not converge. ‘Female EPP’ decreased with the duration of the pair bond (measured until the start of laying), was higher when the sex-ratio was female-biased (only relevant for data from 2005 and 2006), and was not influenced by the inbreeding coefficient of the social partner (Table S2). The random effects ‘Female ID’, ‘Pair ID’ and ‘clutch ID’ (a clutch was defined as having no laying gaps longer than 4 days) explained 9%, 26% and 37% of the variance, respectively.

##### Female fecundity

‘Female fecundity’ (number of eggs laid per breeding round) was square-root transformed to approach normality. Female fecundity increased with the number of days a female spent in the aviary (mean ± SD = 60 ± 23 days, range 1–112), and decreased with female age (mean ± SD = 735 ± 285 days, range 265–1511 days; Table S2). The random effects ‘Female ID’ and ‘Experiment ID’ (18 levels after differentiating testing cohorts and breeding rounds) explained 45% and 10% of the variance, respectively.

#### c) Quantitative Genetic Analyses

We used animal models to carry out quantitative genetic analyses, closely following the initial study [29]. To calculate the parameters, we implemented both a restricted maximum likelihood (REML) method using VCE 6.0.2 [42], and a Bayesian approach using a Monte Carlo-Markov Chain (MCMC) with the package MCMCglmm in R 3.4.0 [48]. Within each type of model (VCE or MCMCglmm), we used two units of analysis: raw data representing single observations and individual mean trait estimates based on the best linear unbiased predictions (BLUPs).

To test the ‘intersexual pleiotropy’ hypothesis, we used four versions of animal models (as in [29]) to estimate the heritability and genetic correlations between aspects of male and female extra-pair mating behavior (five traits: ‘male courtship rate’, ‘male EPP’, ‘female unpaired response’, ‘female extra-pair response’ and ‘female EPP’): a permanent-environment model with repeated measures on individuals in VCE (model 1) and in MCMCglmm (model 2); a model on individual estimates in VCE (model 3) and in MCMCglmm (model 4). For models 3 and 4, individual estimates were BLUPs extracted from the mixed-effect models shown in Table S2. All models are based on the joint data from the initial study [29] and the selection lines.

For comparison between earlier and new findings, we also ran models 1 and 2 on the respective subsets of data (initial data: models 5 and 6 which are updated for model structure compared to the ones published previously; new data: models 7 and 8).

To test the ‘intrasexual pleiotropy’ hypothesis, we used four versions of animal models (similar to models 1 to 4 above) to estimate the heritability and genetic correlations within females (five traits: ‘female fecundity’, ‘female unpaired response’, ‘female extra-pair response’, ‘female within-pair response’ and ‘female EPP’): a permanent-environment model in VCE (model 9) and in MCMCglmm (model 10); a model on individual estimates in VCE (model 11) and in MCMCglmm (model 12). All models are based on the joint data.

